# Genetic insights into the management and conservation of Arctic charr in North Wales

**DOI:** 10.1101/2021.12.03.470319

**Authors:** Samantha V. Beck, Gary R. Carvalho, Ian McCarthy, Walter Hanks, Robert Evans, Robert Edwards, Martin Taylor, Mark de Bruyn

## Abstract

Aquatic species throughout the world are threatened by extinction in many parts of their range, particularly in their most southerly distributions. Arctic charr (*Salvelinus alpinus*) is a Holarctic species with a distribution that includes the glacial lakes of North Wales, towards it southern limit. To date, no genetic studies have been conducted to determine the genetic health of the three remaining native Arctic charr populations in North Wales, despite exposure to stocking and adverse environmental and ecological conditions. We used seven microsatellite loci to determine whether: 1) genetic differentiation existed between native populations; 2) translocated populations from Llyn Peris were genetically similar to the historically connected Llyn Padarn population; and 3) hatchery supplementation negatively impacted genetic diversity in Llyn Padarn. All three native populations retained their genetic integrity, with Llyn Bodlyn showing high levels of divergence (F_ST_ = 0.26 ± 0.02SD) as well as low genetic diversity (H_O_ 0.30) compared to remaining populations (H_O_ 0.64 ± 0.14SD). Although evidence suggests that stocking increased the effective population size of Llyn Padarn in the short term without impacting genetic diversity, the long term effects of such practices are yet to be seen. Results provide baseline data for conservation management, and highlight the need for protection of small isolated populations that are being negatively impacted by the processes of genetic drift due to escalating anthropogenic pressures. Continual monitoring of both Arctic charr and their habitats using a combination of methods will increase the likelihood that these threatened and iconic populations will persist in the future.

## Introduction

Earth’s rich biota is in dramatic decline, with human-induced environmental change being one of the most significant causes of species extinctions. Biodiversity loss is mostly centred around numbers of described species, which may misrepresent the severity of such declines. When focussing on the number of populations, however, results from terrestrial ecosystems show that population extinctions are much greater than species extinctions (Ceballos et al. 2017), suggesting that we have already entered the sixth mass extinction. Freshwater ecosystems are comparatively less well studied. Freshwaters comprise only 0.01% of the Earths’ water, covering just 2.3% of terrestrial areas, yet harbour 9.5% of all known animal species globally, including one-third of vertebrates (Balian et al. 2008). These remarkably rich ecosystems are experiencing declines in biodiversity that are far greater than those of either terrestrial or marine ecosystems (WWF, 2016). Indeed, the IUCN red list have recently assessed half of the worlds’ known freshwater fish species, one third of which were categorised as threatened with extinction (IUCN 2019).

Environmental stressors such as pollution, temperature change, habitat destruction, and overexploitation can result in small isolated populations that are increasingly subject to genetic drift and inbreeding due to declining effective population sizes, resulting in a loss of genetic variation and a decrease in fitness (Hauser et al. 2002; Nowak et al. 2009; Ursenbacher et al. 2009; Gomaa et al. 2011). For example, the construction of hydropower stations have removed spawning sites and migratory routes for various salmonid populations that rely heavily upon freshwater connectivity, due to their high natal philopatry (Mathers et al. 2002; Maitland et al. 2007). Translocations of entire populations across landscapes are often used as a tool to mitigate various anthropogenic stressors and maintain biodiversity (genetic diversity), or increase the number or size of populations under threat (Weeks et al. 2011). However, the movement of populations from native to non-native habitats can often result in low fitness levels due to the disruption of local adaptation, thus having a major impact on the genetic composition and evolutionary trajectories on wild populations (Allendorf et al. 2008; Valiquette et al. 2014).

Stocking practices are another important conservation management tool that aims to alleviate such genetic erosion in fishes, yet the introduction of foreign genetic material, i.e. through the release of translocated and/or hatchery-reared individuals, may negatively impact the evolutionary potential and genetic integrity of populations (Araki et al. 2008; Laikre et al. 2010; Perrier et al. 2013; Quiñones et al. 2014; Valiquette et al. 2014), even within a single generation (Araki et al. 2007a; Christie et al. 2012b). Salmonids are some of the most intensively propagated species in hatchery stocking programs (Lackey et al. 2006), due in part to their socio-economic importance. Despite the importance of intraspecific diversity for a species’ resilience to environmental change (Sgrò et al. 2011; Pauls et al. 2013), many salmonid populations across their native range in the northern hemisphere are experiencing unprecedented declines (Winfield et al. 2010; ICES 2019). The capability of hatchery programmes in maintaining both genetic diversity and fitness in the wild over the long term is uncertain (Fraser 2008; Christie et al. 2012a).

The Holarctic distribution of Arctic charr makes this salmonid species particularly vulnerable to rising temperatures and increasing anthropogenic pressures, especially at their southern extremes (Kovach et al. 2019; Muhlfeld et al. 2019). Numerous translocations and hatchery supplementations of Arctic charr (‘Torgoch’) in North Wales have occurred as a response to the negative impacts these environmental and anthropogenic pressures have had on natural charr populations. There were originally four known native populations of Arctic charr in North Wales: Llyn(-lake) Padarn, Llyn Peris, Llyn Cwellyn and Llyn Bodlyn (Fig. 1). However, the construction of the Dinorwig power station in 1974 saw the translocation of Arctic charr from Llyn Peris (the type locality for *Salvelinus alpinus perissi*; Günther, 1862) to Ffynnon Llugwy and Llyn Dulyn after allozyme evidence suggested no genetic differentiation between Arctic charr from Llyn Peris and those from Llyn Padarn (Child 1977). Not only did this construction result in the extirpation of Llyn Peris charr, but also removed spawning habitat for Arctic charr in Llyn Padarn, which were believed to migrate into Llyn Peris to spawn (Butterworth 1980). Given the high productivity of Llyn Padarn and the propensity of Arctic charr to diverge along trophic resource gradients (Skúlason et al. 1999), as well as evidence that both populations differed in size and parasite abundance (McCarthy 2007), it is likely that Arctic charr in Llyn Padarn and Llyn Peris exhibited, to some extent, parapatric divergence both phenotypically (McCarthy 2007) and genetically. Since the construction of Dinorwig power station, numerous translocations of Arctic charr to lakes that are deeper and at higher altitudes have been conducted across North Wales, where there now exist not only three native populations, but also six translocated populations (see reviews by (Maitland et al. 2007; McCarthy 2007; Ferguson et al. 2019). Although Child (1977) found genetic differentiation between the native populations (Llyn Padarn/Peris, Llyn Cwellyn and Llyn Bodlyn), no genetic study using DNA-based molecular markers has since been conducted on these Welsh Arctic charr populations, and their status remains unknown. In Llyn Padarn, for example, a sharp decline of Arctic charr in 2009 due to a toxic algal bloom (amongst various other anthropogenic stressors; (White 2012; Clabburn et al. 2014; Hatton-Ellis 2016) resulted in the establishment of a back-up population in Llyn Crafnant, from a broodstock created from a total of 77 females and 54 males (from 2009-2013; Table 1) from Llyn Padarn for use in a hatchery supplementation programme. However, the effects of such stocking practices remain unclear and may potentially have a negative impact on the wild population (Araki and Schmid 2010).

**Figure 1.**
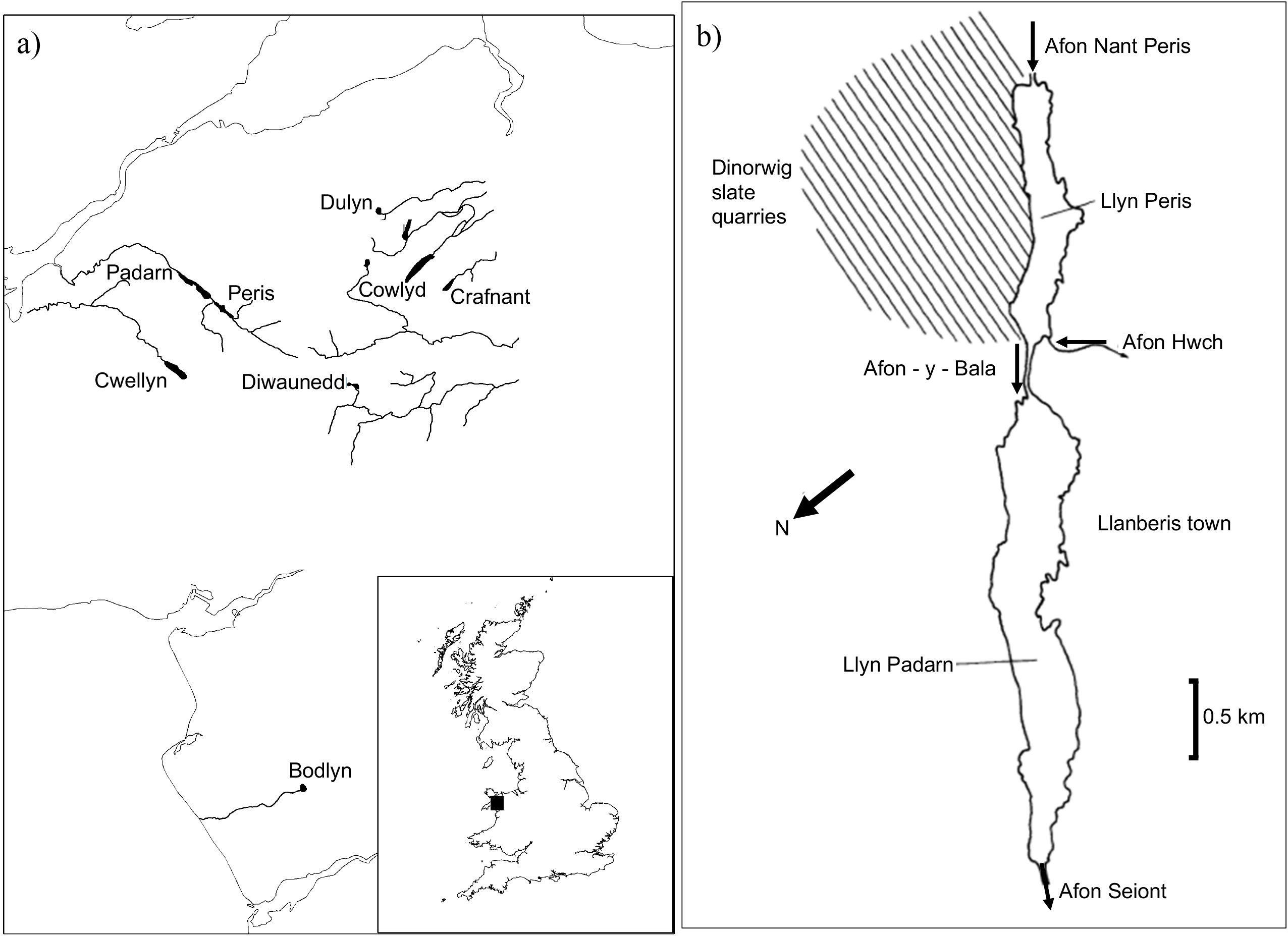
Maps of a) the distribution of native (Padarn, Peris, Bodlyn and Cwellyn) and translocated (Dulyn, Ffynnon Llugwy, Cowlyd and Diwaunedd) Arctic charr populations in North Wales includeed in this study, modified from McCarthy (2007); and b) the historical connection between Llyn peris and Llyn Padarn before the construction of a hydropower dam, modified from (Elner et al. 1980).

**Table 1.**
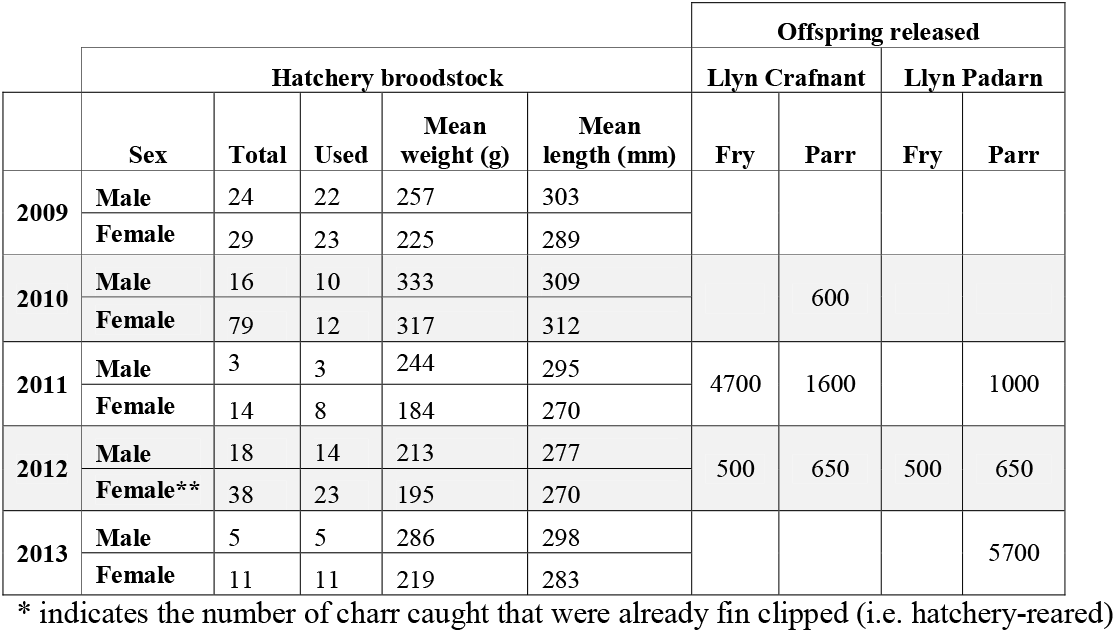
A summary of the number of adult Arctic charr (*Salvelinus alpinus*) caught and used as hatchery broodstock, as well as the number of offspring successfully released into Llyn Crafnant as a back-up population, and Llyn Padarn.

|  | Hatchery broodstock |  |  |  |  | Offspring released |  |  |  |
| --- | --- | --- | --- | --- | --- | --- | --- | --- | --- |
|  |  |  |  |  |  | Llyn Crafnant |  | Llyn Padarn |  |
|  | Sex | Total | Used | Mean weight (g) | Mean length (mm) | Fry | Parr | Fry | Parr |
| 2009 | Male | 24 | 22 | 257 | 303 |  |  |  |  |
|  | Female | 29 | 23 | 225 | 289 |  |  |  |  |
| 2010 | Male | 16 | 10 | 333 | 309 |  | 600 |  |  |
|  | Female | 79 | 12 | 317 | 312 |  |  |  |  |
| 2011 | Male | 3 | 3 | 244 | 295 | 4700 | 1600 |  | 1000 |
|  | Female | 14 | 8 | 184 | 270 |  |  |  |  |
| 2012 | Male | 18 | 14 | 213 | 277 | 500 | 650 | 500 | 650 |
|  | Female** | 38 | 23 | 195 | 270 |  |  |  |  |
| 2013 | Male | 5 | 5 | 286 | 298 |  |  |  | 5700 |
|  | Female | 11 | 11 | 219 | 283 |  |  |  |  |

\* indicates the number of charr caught that were already fin clipped (i.e. hatchery-reared)

This study aims to understand how both historic and contemporary management practices have influenced the genetic variability of small freshwater populations. Using seven microsatellite markers, we specifically asked whether: 1) genetic differentiation and genetic integrity exists between all three remaining native populations of Arctic charr in North Wales; 2) translocated Arctic charr from Llyn Peris were genetically similar to Arctic charr in Llyn Padarn; and 3) stocking of hatchery-reared charr reduced the genetic variability of the wild population in Llyn Padarn.

## Methods

### Sample collection

A total of 307 individuals (fin clips) from eight lakes in North Wales (Fig. 1) were used in this study (see Table 2 for lake information and Table 3 for sample sizes). Fyke nets were placed near the entrance to the Afon-y-Bala (Fig. 1b) in Llyn Padarn as part of Natural Resources Wales (NRW) hatchery-supplementation programme. Fin clips were taken and stored in 96% ethanol from each individual after stripping had occurred. The fertilised eggs were then reared in Mawddach hatchery and subsequently released into Llyn Crafnant as a secondary population for Llyn Padarn (Table 1). All other populations were also sampled using fyke nets by McCarthy in 2008 and stored in 96% ethanol, until DNA extractions were conducted in 2013.

**Table 2.** The number of known Arctic charr populations in North Wales and their status’, modified from McCarthy (2007).

| Lake | NGR | Altitude (m) | Surface area (ha) | Max. depth (m) | Use of waterbody | Date of transfer | Status | Other salmonid species |
| --- | --- | --- | --- | --- | --- | --- | --- | --- |
| **Native populations** |  |  |  |  |  |  |  |  |
| Llyn Padarn<sup>a</sup> | SH572613 | 108 | 102 | 29 | Recreation |  | Declining | *S. trutta*, *S. salar* |
| Llyn Peris | SH591594 | 111 | 37 | 34 | Pump storage reservoir |  | Extinct |  |
| Llyn Cwellyn<sup>a</sup> | SH560549 | 146 | 89 | 40 | Water supply |  | Declining | *S. trutta*, *S. salar* |
| Bodlyn<sup>a</sup> | SH648239 | 385 | 16 | 22 | Water supply |  | Unknown | *S. trutta* |
| **Translocated populations** |  |  |  |  |  |  |  |  |
| Ffynnon Llugwy<sup>a</sup> | SH692627 | 550 | 15 | 45 | Reservoir | 1976-1981 | Resident, healthy | *S. trutta* |
| Llyn Dulyn<sup>a</sup> | SH700665 | 538 | 13 | 58 | Reservoir | 1982-1985 | Resident, healthy | *S. trutta* |
| Llyn Cowlyd<sup>a</sup> | SH727623 | 370 | 116 | 68 | Reservoir |  | Resident, healthy | *S. trutta* |
| Llynau Diwaunedd<sup>a</sup> | SH682537 | 371 | 7 & 5 |  | None |  | Resident, unknown | *S. trutta* |
| Llyn Coedty | SH754666 | 277 | 5 |  | Reservoir |  | Transient | *S. trutta* |
| Llyn Eigiau | SH720652 | 375 | 46 | 10 | Reservoir |  | Transient | *S. trutta* |
| Llyn Crafnant<sup>a</sup> | SH750611 | 183 | 25 | 22 | Reservoir |  | Hatchery stock from Llyn Padarn | *S. trutta* |

<sup>a</sup> Populations included in this study
NGR, National Grid Reference

**Table 3.**
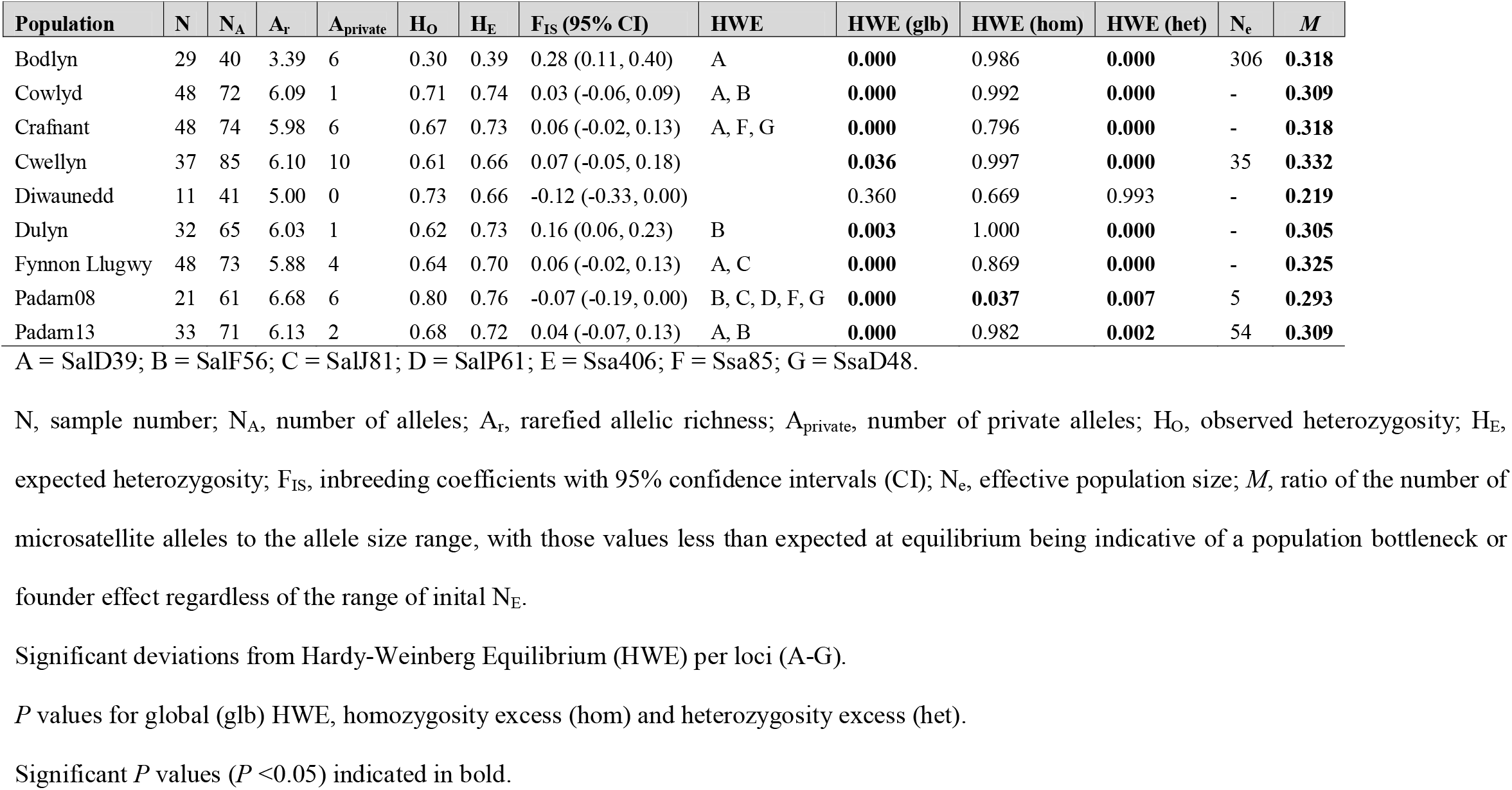
Genetic diversity indices of eight populations of Arctic charr (*Salvelinus alpinus*) in North Wales. Effective population sizes (N_e_) are n for all native populations, including before (Padarn08) and after (Padarn13) stocking in Llyn Padarn.

| Population | N | N<sub>A</sub> | A<sub>r</sub> | A<sub>private</sub> | H<sub>O</sub> | H<sub>E</sub> | F<sub>IS</sub> (95% CI) | HWE | HWE (glb) | HWE (hom) | HWE (het) | N<sub>e</sub> | M |
| --- | --- | --- | --- | --- | --- | --- | --- | --- | --- | --- | --- | --- | --- |
| Bodlyn | 29 | 40 | 3.39 | 6 | 0.30 | 0.39 | 0.28 (0.11, 0.40) | A | **0.000** | 0.986 | **0.000** | 306 | **0.318** |
| Cowlyd | 48 | 72 | 6.09 | 1 | 0.71 | 0.74 | 0.03 (-0.06, 0.09) | A, B | **0.000** | 0.992 | **0.000** | - | **0.309** |
| Crafnant | 48 | 74 | 5.98 | 6 | 0.67 | 0.73 | 0.06 (-0.02, 0.13) | A, F, G | **0.000** | 0.796 | **0.000** | - | **0.318** |
| Cwellyn | 37 | 85 | 6.10 | 10 | 0.61 | 0.66 | 0.07 (-0.05, 0.18) |  | **0.036** | 0.997 | **0.000** | 35 | **0.332** |
| Diwaunedd | 11 | 41 | 5.00 | 0 | 0.73 | 0.66 | -0.12 (-0.33, 0.00) |  | 0.360 | 0.669 | 0.993 | - | **0.219** |
| Dulyn | 32 | 65 | 6.03 | 1 | 0.62 | 0.73 | 0.16 (0.06, 0.23) | B | **0.003** | 1.000 | **0.000** | - | **0.305** |
| Fynnon Llugwy | 48 | 73 | 5.88 | 4 | 0.64 | 0.70 | 0.06 (-0.02, 0.13) | A, C | **0.000** | 0.869 | **0.000** | - | **0.325** |
| Padarn08 | 21 | 61 | 6.68 | 6 | 0.80 | 0.76 | -0.07 (-0.19, 0.00) | B, C, D, F, G | **0.000** | **0.037** | **0.007** | 5 | **0.293** |
| Padarn13 | 33 | 71 | 6.13 | 2 | 0.68 | 0.72 | 0.04 (-0.07, 0.13) | A, B | **0.000** | 0.982 | **0.002** | 54 | **0.309** |

A = SalD39; B = SalF56; C = SalJ81; D = SalP61; E = Ssa406; F = Ssa85; G = SsaD48.
N, sample number; N<sub>A</sub>, number of alleles; A<sub>r</sub>, rarefied allelic richness; A<sub>private</sub>, number of private alleles; H<sub>O</sub>, observed heterozygosity; H<sub>E</sub>, expected heterozygosity; F<sub>IS</sub>, inbreeding coefficients with 95% confidence intervals (CI); N<sub>e</sub>, effective population size; M, ratio of the number of microsatellite alleles to the allele size range, with those values less than expected at equilibrium being indicative of a population bottleneck or founder effect regardless of the range of initial N<sub>E</sub>.
Significant deviations from Hardy-Weinberg Equilibrium (HWE) per loci (A-G).
*P* values for global (glb) HWE, homozygosity excess (hom) and heterozygosity excess (het).
Significant *P* values (*P* < 0.05) indicated in bold.

### DNA extractions

DNA was extracted from adipose fin clippings for all 307 individuals using the CTAB (hexadecyltrimethyammonium bromide) extraction method. Samples were suspended in CTAB extraction buffer: 100mM Tris-HCl pH 8.0; 1.4M NaCl; 20mM EDTA pH 8.0; 2% CTAB; 0.2% 2-mercaptoethanol, and 0.1mg/ml of proteinase K was added to digest the samples overnight at 60°C. Two chloroform:isoamylalcohol washes were then performed followed by sodium acetate precipitation (3M, pH4.8), and one wash of 70% ethanol before DNA extracts were suspended in ultrapure H_2_0 and stored at −20°C.

### DNA amplification and genotyping

All 307 individuals were analysed at eight microsatellite loci using QiagenType-it Multiplex PCR kit. Each 10μl multiplex reaction contained at least 2ng/μl DNA template, 2.5μl QiagenType-it Multiplex reaction buffer, 50mg/ml Bovine Serum Albumin (BSA) due to potential degradation of DNA, and primers at different concentrations (Table S1). PCR conditions were the same for both multiplexes, and began with an initial 95°C activation step, followed by 34 cycles of 30s at 94°C, 90s at 57°C, and 60s at 72°C, followed by 31 cycles of 30s at 94°C, 90s at 50°C, and 60s at 72°C, and finalised with a 30min extension phase at 60°C. A subset of amplified products were migrated via electrophoresis, and then all samples were run on the Applied Biosystems ABI 3130xl Genetic Analyser, and microsatellite allele sizes scored using GENEMAPPER 3.0 (Applied Biosystems).

### Genetic diversity

Unless otherwise stated, all analyses were conducted in R (R Core Team 2016). The presence of null alleles were determined using the package *PopGenReport* v3.0.4 (Adamack and Gruber 2014), as well as the identification of any private alleles (A_private_). For each individual locus, observed (H_O_) and expected (H_E_) heterozygosities were calculated using the *adegenet* package (Jombart 2008), whilst *hierfstat* (Goudet 2005) and *PopGenReport* (Adamack and Gruber 2014) was used to calculate inbreeding coefficients (F_IS_) with bootstrapping over loci to provide confidence intervals for each population, and rarified allelic richness (A_r_), respectively. The package *pegas* (Paradis 2010) was used to test for deviations from Hardy-Weinberg equilibrium (HWE) for each locus. The *diveRsity* package (Keenan et al. 2013) was used to calculate several genetic parameters across loci: A_r_ with rarefaction; F_IS_ using 10,000 bootstrap replicates across individuals were used to generate 95% confidence intervals; H_O_ and H_E_; global HWE(glb) based on log-likelihood tests for goodness of fit using 10,000 Monte Carlo replicates; one-tailed tests for heterozygosity deficiency (HWE(hom)) and heterozygosity excess (HWE(het)).

### Population structure

The extent of genetic differentiation between populations was quantified using Nei’s pairwise F_ST_ (Nei 1973) in the package *hierfstat* (Goudet 2005), with 10,000 bootstraps over loci of pairwise F_ST_ to determine confidence intervals. Hierarchical analyses of molecular variance (AMOVA; Excoffier et al., 1992) was used to determine the source of genetic differentiation, using the package *ade4* (Dray and Dufour 2007). This approach partitions variance into covariance components to calculate Phi’s ‘Φ’ fixation indices for various hierarchical levels. We conducted two AMOVAs, the first of which had two hierarchical levels: 1) populations, and 2) stocking status (e.g. stocked versus unstocked). Neither the Crafnant hatchery population, nor the Padarn population after stocking were included in this part of the analyses as their potential genetic similarities were likely to bias results. The second AMOVA used data from Llyn Padarn only to test for differences in variation before and after stocking. Significance was tested using 1,000 permutations.

The Bayesian clustering programme STRUCTURE (Pritchard et al. 2000) was implemented within *stratag* (Archer et al. 2017) without admixture due to the landlocked status of each population and the lack of resident Arctic charr in those translocated lakes. A burn-in period was set to 100,000 and 100,000 Markov Chain Monte Carlo (MCMC) iterations, with *k* ranging from 2–9. The best *k* was chosen according to Δk (Evanno et al. 2005) and *ggplot* (Wickham 2016) was used to plot STRUCTURE outputs. Finally, Discriminate Analysis of Principal Components (DAPC) were performed within *adegenet* (Jombart 2008) to describe the relationships between each genetic cluster at two levels: 1) between all clusters identified from STRUCTURE; and 2) between the hatchery stock (Crafnant) and the Padarn populations before (Padarn08) and after (Padarn13) hatchery supplementation.

### Effective population size and bottlenecks

Effective population size (N_e_) was estimated using the linkage disequilibrium method in native populations only, due to the potential bias caused by population admixture in those translocated populations (Araki et al. 2007b; Waples and Do 2010). Such estimates were implemented using the linkage disequilibrium method within the software NeEstimator v2.0 (Do et al. 2014). In a population that has reduced in size, genetic drift is more likely to have resulted in the loss of rare alleles than common alleles. As such, the ratio of microsatellite allele number to microsatellite allele size range (*M*; Garza and Williamson, 2001) is expected to be reduced following a population bottleneck. We therefore used the programme M_P_Val (Garza and Williamson, 2001) to determine whether populations have experienced a recent bottleneck, evident through smaller *M* in relation to equilibrium expectations. Using recommended paramenters by Garza and Williamson (2001), we used a two-phase mutation model with 90% single-step mutations, with the average size of multi-step mutations being 3.5 repeat units. We assumed a microsatellite mutation rate of 5 × 10^−4^ (Garza and Williamson 2001) and removed all monomorphic loci from the analysis. The N_e_ of populations before any reduction in size is unkown, therefore the analysis was run using a range of suitable N_e_ estimates: N_e_ = 250 (Θ = 0.5); N_e_ = 500 (Θ = 1); and N_e_ = 1000 (Θ = 2).

Evidence of a population decline was indicated if <5% of the 10,000 simulated replicates fell below the critical M value (*M*_c_), calculated using Critical_M (Garza and Williamson 2001).

## Results

### Genetic diversity

The only locus that was removed due to the presence of null alleles was SalE38. Although Crafnant had population-specific null alleles for SalD39 and Ssa406, their removal did not change the results and they were therefore retained within the analyses, leaving a total of seven microsatellite loci. The total number of alleles observed per locus ranged from seven (SalF56) to 35 (SsaD48), with a mean of 18.14 (Table S2). Bodlyn exhibited fixed alleles at two loci: SalF56 and SalJ81, and also had the lowest mean A_r_ (3.39) across all populations, with Padarn08 having the highest mean A_r_ (6.68), with mean rarified A_R_ of 5.70 (±0.97, SD). Such patterns were also reflected in expected levels of heterozygosity (Table 3), with Bodlyn having the lowest H_e_ (0.39) and Padarn08 having the highest H_e_ (0.76) levels compared to the overall average of 0.68 (±0.11, SD). Bodlyn also shows the highest F_IS_ values (0.28; albeit not significant).

Of the nine populations examined, eight exhibited significant deviations from HWE. Only Diwaunedd showed no deviations from HWE, but this population also had the lowest sample size (N=11). Seven of the remaining populations deviated from HWE as a result of an excess of heterozygosity (Bodlyn, Cowlyd, Crafnant, Cwellyn, Dulyn, Fynnon Llugwy and Padarn13), whilst Padarn08 deviated from HWE as a result of heterozygote deficiency as well as heterozygote excess (Table 1).

### Population structure

Results showed the native Bodlyn population to be highly differentiated from all other populations (Table 4). Estimates of global F_ST_ amongst all nine populations was 0.16, with pairwise differences between Llyn Bodlyn and all other populations averaging 0.26 (±0.02, SD). A translocated population, Llyn Dulyn, also showed significant genetic differentiation between all populations except Diwaunedd and Padarn08, with an average F_ST_ of 0.07 (±0.10, SD). There was no genetic differentiation between Padarn before (Padarn08) or after (Padarn13) stocking, nor was there any differences compared with the hatchery population (Crafnant). Excluding Llyn Crafnant and Padarn13, AMOVAs revealed that genetic differentiation between the stocked (i.e. translocated) versus unstocked populations (i.e. native) explained 6.87% of the total variance, populations explained 14.88%, and between individuals within a population and within individuals explained 8.22% and 70.03%, respectively (all *P* <0.05; Table 5). When examining the effects of hatchery supplementation on Llyn Padarn charr, AMOVAs found 2.96% (*P* <0.001) differences in genetic differentiation before and after supplementation (F_ST_) (Table 5; Fig. S1). Whilst variation between individuals within populations (F_IS_), and variation within individuals (F_IT_) showed no differences (Table 5).

**Table 4.** Pairwise F_ST_ between all pairs of Arctic charr populations in North Wales, with those significant values (P<0.05) in bold.

|  | Bodlyn | Cowlyd | Crafnant | Cwellyn | Diwaunedd | Dulyn | Ffynnon Llugwy | Padarn08 |
| --- | --- | --- | --- | --- | --- | --- | --- | --- |
| Cowlyd | **0.247** |  |  |  |  |  |  |  |
| Crafnant | **0.228** | 0.034 |  |  |  |  |  |  |
| Cwellyn | **0.262** | 0.107 | 0.122 |  |  |  |  |  |
| Diwaunedd | **0.281** | 0.041 | 0.054 | 0.047 |  |  |  |  |
| Dulyn | **0.271** | **0.015** | **0.035** | **0.124** | 0.063 |  |  |  |
| Ffynnon Llugwy | **0.267** | **0.011** | 0.040 | 0.116 | 0.043 | **0.014** |  |  |
| Padarn08 | **0.269** | 0.021 | 0.031 | 0.120 | 0.085 | 0.018 | 0.025 |  |
| Padarn13 | **0.275** | **0.014** | 0.019 | 0.118 | 0.059 | **0.018** | **0.016** | 0.023 |

**Table 5.** Analysis of molecular variance (AMOVA) to detect population differentiation in Arctic charr in North Wales for: 1) Llyn Padarn before and after stocking, and 2) all stocked lakes versus unstocked lakes (excluding Crafnant and Padarn13). Significant differences in bold.

| Source of variation | DF | SS | MS | Sigma | Variation (%) | Phi | P |
| --- | --- | --- | --- | --- | --- | --- | --- |
| Analysis of Padarn before and after stocking |  |  |  |  |  |  |  |
| **Between stocks (before and after stocking)** | **1** | **13.61** | **13.61** | **0.16** | **2.96** | **0.03** | **0.001** |
| Between individuals within Stocked | 52 | 280.78 | 5.40 | 0.16 | 3.02 | 0.03 | 0.169 |
| Within individuals | 54 | 274 | 5.07 | 5.07 | 94.02 | 0.06 | 0.065 |
| Total | 107 | 568.39 | 5.31 | 5.40 | 100 |  |  |
| Analysis of stocked versus unstocked lakes (no Crafnant/Padarn13) |  |  |  |  |  |  |  |
| **Between stocked and unstocked** | **1** | **166.14** | **166.1** | **0.44** | **6.87** | **0.07** | **0.032** |
| **Between populations within stocked/unstocked** | **5** | **322.09** | **64.42** | **0.95** | **14.88** | **0.16** | **0.001** |
| **Between individuals within population** | **219** | **1214.83** | **5.547** | **0.53** | **8.22** | **0.11** | **0.001** |
| **Within individuals** | **226** | **1015.38** | **4.493** | **4.49** | **70.03** | **0.30** | **0.001** |
| Total | 451 | 2718.43 | 6.028 | 6.42 | 100.00 |  |  |

DF, degrees of freedom; SS, sum of squares; MS, mean squares

For STRUCTURE analysis, the most likely number of clusters identified by Δk was *k* = 3. Each of these three genetic clusters corresponded to one of the three remaining native populations: Bodlyn, Padarn and Cwellyn (Fig. 2a). All three native populations seem to have conserved their genetic integrity, as shown by both DAPC and STRUCTURE plots (Fig. 2). The Diwaunedd stock seems to have primarily originated from Llyn Cwellyn (Fig. 2a), but the small sample size (N = 11) likely prevents its clear clustering, and it is instead situated inbetween Padarn and Cwellyn (Fig. 2b). Although the hatchery Crafnant population primarily consists of genotypes from Llyn Padarn, there also appears to be some genotypes from Llyn Cwellyn as well (Fig. 2a). However, such patterns were not reflected within the DAPC (Fig. 2b).

**Figure 2.**
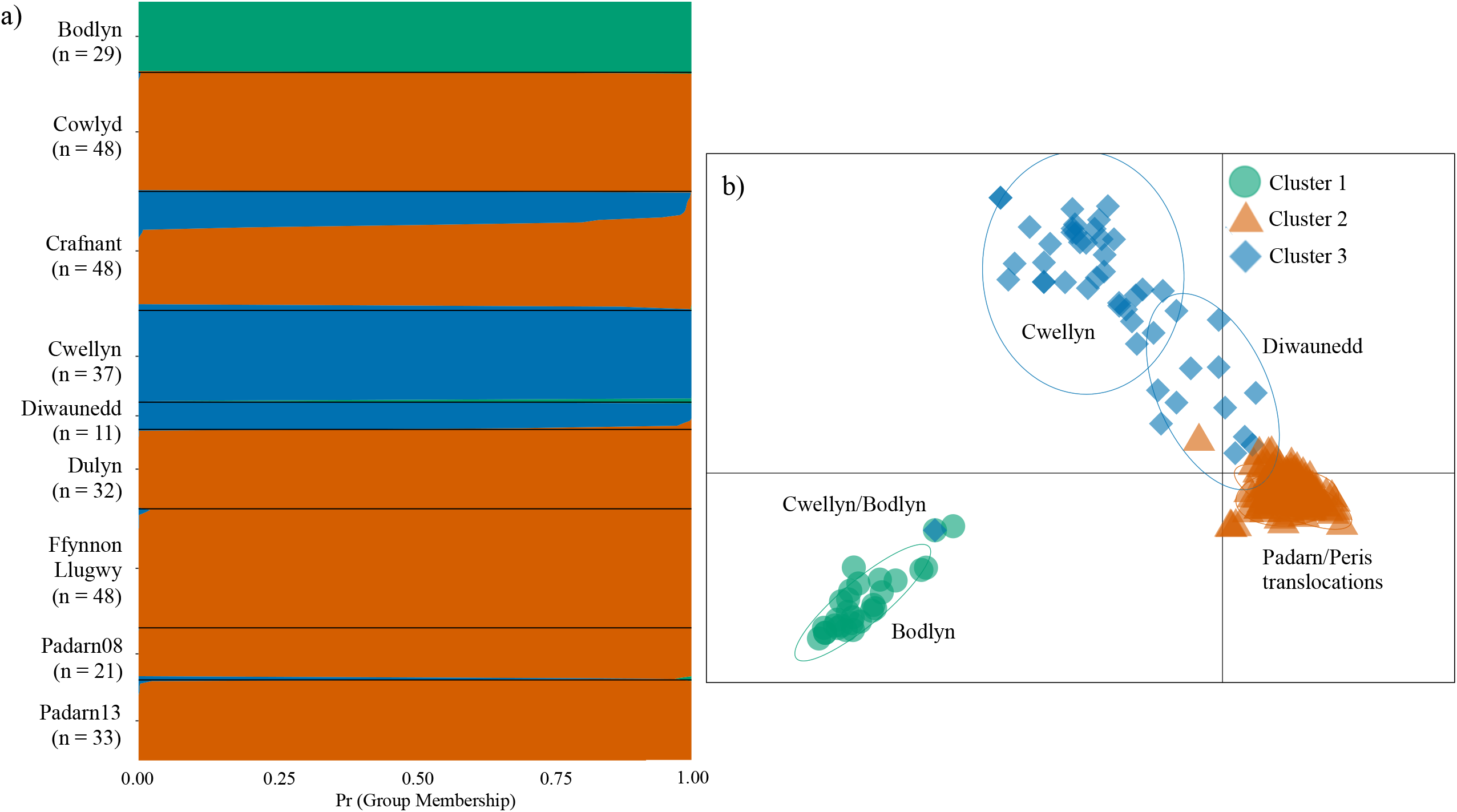
Results from a) STRUCTURE and b) Discriminant Analysis of Principle Components (DAPC) demonstrating the genetic groupings of genotypes according to the most likely number of clusters determined by Δ *k*. Symbols correspond to different clusters (i.e. genetic group).

### Estimates of effective population size and bottlenecks

Estimates of N_e_ for native populations varied from as little as five for Padarn08 (before hatchery supplementation), to 306 for Llyn Bodlyn. Llyn Cwellyn had an N_e_ of 35, whilst Padarn13 (after hatchery supplementation) was estimated at 54 (Table 3). All populations, both native and translocated, show evidence of a population bottleneck or founder effects, respectively (*P* <0.001), with *M* ratios (*M*: 0.219 – 0.332) showing a lower value than expected under equilibrium conditions (*M*_c_: 0.696 - 0.718), regardless of N_e_ before population size reduction (Table 3).

## Discussion

Despite stocking practices and numerous translocations of Arctic charr between lakes in North Wales, the population genetic structure amongst the three remaining native populations has been retained. Hatchery supplementation of the declining native Llyn Padarn population has increased effective population size (N_e_) in the short term, but at the cost of a reduction in the number of private alleles, i.e. loss of genetic uniqueness (Table 4). All three native populations have retained their genetic integrity, yet the isolated location of Llyn Bodlyn has likely exposed this population to genetic drift, evident through low levels of genetic diversity. All populations show evidence for a recent population bottleneck or, for those translocated populations, founder effects. This study highlights the need to further conserve the genetic diversity of Arctic charr in North Wales, especially in Llyn Bodlyn where there is currently no conservation protection, despite harbouring a highly divergent and vulnerable native population of Arctic charr.

### Genetic diversity in an isolated population

All populations of Arctic charr in North Wales – with the exception of Llyn Bodlyn (H_O_ = 0.30; A_R_ = 3.39) – showed moderate-high levels of genetic diversity (H_O_ = 0.68 ± 0.06; A_R_ = 5.99 ± 0.47; Table 3) compared to other British Arctic charr populations (H_O_ = 0.58 ± 0.15 SD; A_R_ = 6.52 ± 2.51 SD; Wilson et al., 2004). The high genetic differentiation of Llyn Bodlyn (F_ST_ = 0.26 ± 0.02; *P* <0.05) reflects its isolated position (∼50km from the nearest population) as well as its relatively high elevation (385m), which is three-times higher than all other native populations (Table 2). Although such elevation might buffer against increasing temperatures (Roberts et al. 2013), Llyn Bodlyn is the smallest (16ha) and shallowest (22m) of all Arctic charr lakes in North Wales. Furthermore, this lake is also a reservoir that can have up to 60% of its storage drained in periods of drought (plus more in severe cases; (DCWW 2015)), leaving little suitable habitat for Arctic charr during hot summer periods. These small isolated populations are most vulnerable to the negative impacts of genetic drift, reducing heterozygosity and increasing levels of inbreeding (Reed and Frankham 2003), especially when exposed to such stochastic environmental changes. Here we suggest that Llyn Bodlyn has been subject to such processes, evident from the low levels of heterozygosity (H_O_ = 0.30), allelic richness (A_R_ = 3.39) and high inbreeding coefficient (F_IS_ = 0.28) compared to the overall mean for all populations (H_O_ = 0.64 ± 0.14 SD; A_R_ = 5.70 ± 0.97 SD; and F_IS_ = 0.10 ± 0.09 SD; see Table 3). Nevertheless, estimates of N_e_ are high (307) compared to other native populations such as Llyn Cwellyn (N_e_ = 35), suggesting that habitat limitations on population size have, over a long period of time, reduced the genetic variability of this isolated population (Franklin 1980). It is important to note that whilst estimates of N_e_ are a valuable measure for conservation management, assumptions around N_e_ values are often violated in nature (i.e. no selection, migration, mutation and discrete generations). Almost all populations in this study deviate from Hardy-Weinberg equilibrium (HWE), and as such, estimates of N_e_ should be approached with caution.

### Genetic integrity

All three native populations of Arctic charr retained their genetic differentiation and integrity (Fig. 2; Table 4), with Llyn Cwellyn (A_private_ = 10) having almost twice the number of private alleles compared to Llyn Bodlyn and Llyn Padarn before stocking (A_private_ = 6). Before the homogenising effects of stocking, Llyn Padarn (Padarn08) showed the highest levels of genetic variability (H_O_ = 0.80; A_r_ = 6.68), but also has an alarmingly low estimate of N_e_ (N_e_ = 5). Such results are indicative of a recent population bottleneck, which is often accompanied by a temporary increase in heterozygosity (Garza and Williamson 2001). Indeed, all three native populations showed evidence of a recent bottleneck (*M* = 0.293-0.332), with Llyn Padarn prior to stocking having one of the lowest estimates of *M* (0.293) compared to what is expected from this population at equilibrium (0.751). Although hatchery supplementation of Llyn Padarn increased the N_e_ from 5 to 54, such stocking practices may have come at the expense of a reduction in genetic uniqueness, with the number of private alleles reducing from 6 to 2 (Table 2). Results from AMOVA reveal a 2.96% (*P* <0.001) difference in genetic differentiation between Llyn Padarn before compared to after stocking (Fig. S1). However, we found that genetic variability of Llyn Padarn stayed relatively similar to the levels before stocking (H_O_ = 6.68) compared to after stocking (H_O_ = 6.13). Nevertheless, our results suggest that the hatchery stock at Crafnant, which is used to supplement Arctic charr for Llyn Padarn, may contain some Llyn Cwellyn genotypes. Salmonid populations are often repeatedly re-stocked over numerous years, not only increasing the likelihood of populations becoming reliant upon hatchery supplementation and at risk of collapse in the absence of such stocking, but also resulting in reduced recruitment performance of natural populations (Chilcote et al. 2011). For populations to persist in the long term, it is the source of their decline that needs to be addressed before using stock enhancement as a management strategy. However, conservation is also a crisis discipline and such time may rarely be available.

### Lack of historical differentiation

All translocated populations showed evidence of founder effects (Table 3), yet such reductions in population size was not reflected by low levels of genetic diversity (H_O_ = 0.68 ± 0.04 SD). These populations primarily consist of Llyn Padarn genotypes, with the exception of Diwaunedd which contains mostly Llyn Cwellyn genotypes (Fig. 2a). The power of this study, based on a relatively small number of microsatellite markers, might have limited our ability to detect any fine-scale population structure that may have existed between Llyn Padarn and Llyn Peris. Although we lack any genetic evidence for such differentiation, previous studies show phenotypic divergence in body size, with Arctic charr in Llyn Padarn growing larger than those in Llyn Peris due to the increased amount of nutrients in Llyn Padarn (Butterworth 1980; McCarthy 2007). Arctic charr are famed for their phenotypic plasticity, whereby multiple morphs can occur in sympatry, diverging in resources (i.e. habitat and diet) and associated trophic morphologies, body size, behaviour and life-history traits (Skúlason et al. 1996, 1999; Klemetsen 2010; Ferguson et al. 2019). Patterns of phenotypic divergence between Llyn Padarn and Llyn Peris indicates differential habitat use and dietary preferences between the two populations (Butterworth 1980; McCarthy 2007). The consideration of both phenotypic and genetic diversity is important to consider in future management actions, since it is this intraspecific biodiversity that can spread the risk of species losses, as differential responses of locally adapted populations can buffer against environmental perturbations (Figge 2004). The historic dispersal between Llyn Padarn and Llyn Peris may have once buffered the impacts of environmental pressures via metapopulation dynamics (Akçakaya et al. 2007). Although any effects of such loss of gene flow is not reflected by the high levels of genetic diversity, it has likely contributed to the vulnerability of Arctic charr to stochastic environmental events, e.g. the toxic algal bloom in 2009 (BBC 2010).

### Summary

The conservation of genetic diversity is essential to maintain the evolutionary potential of natural populations in response to environmental pressures (Allendorf 2013). Populations in larger habitats are likely to be less susceptible to the loss of such genetic diversity due to bottlenecks because they are not only able to support larger populations, but can also provide more refugial opportunities under adverse conditions (Neville et al. 2009). The increased genetic differentiation in Llyn Bodlyn combined with this populations’ low genetic diversity suggests that genetic drift has been the primary cause of genetic divergence, likely due to the lack of habitat availability - an important consideration for maintaining the evolutionary potential of isolated populations (Whiteley et al. 2010). Whilst hatchery supplementation in Llyn Padarn has been successful at increasing short-term N_e_, achieving a self-sustaining increase in population abundance depends on numerous factors, including whether stocked fish displace naturally produced offspring, as well as the potential negative impact on the fitness of the wild population (Caroffino et al. 2008). The findings from this study highlight the importance of monitoring the few remaining native populations of Arctic charr in North Wales, as well as ensuring that they retain their genetic integrity. Conservation of this genetic diversity is vital for maintaining evolutionary potential, especially in isolated populations with limited habitat availability.

## Acknowledgments

We thank those at Natural Resources Wales and Bangor University who were involved in the collection of samples. Thanks also to the Knowledge Economy Skills Scholarship for funding and providing support for the research.

## Declarations

### Ethics approval

Samples were collected under licenses granted by Natural Resource Wales (NRW; previously the Environment Agency Wales). NRW collected finclips from Arctic charr as part of a normal sampling scheme, a recognised agricultural/husbandry practice performed in accordance with other animal welfare for the purposes of managing and conserving the animals, and have confirmed that this is not a regulated procedure under the Animals (Scientific Procedures) Act.

### Consent to participate

Non applicable

### Consent for publication

Non applicable

### Availability of data and material

Data will be made available on Dryad upon publication of the manuscript

### Code availability

Non applicable

### Authors’ contributions

MT, GRC and MdB conceptualised and designed the study. Samples were collected by WH, IM, RE and RE. Data collection and analysis were performed by SVB. The first draft of the manuscript was written by SVB and all authors commented on subsequent versions of the manuscript. All authors read and approved the final manuscript.

## References

Adamack AT, Gruber B (2014) PopGenReport: simplifying basic population genetic analyses in R. Methods Ecol Evol 5:384–387. https://doi.org/10.1111/2041-210X.12158

Akçakaya HR, Mills M, Doncaster C (2007) The role of metapopulations in conservation. In: Macdonald DW, Service K (eds) Key topics in conservation biology. Blackwell Publishing, Oxford, UK, pp 64–84

Allendorf F, England P, Luikart G, et al (2008) Genetic effects of harvest on wild animal populations. Trends Ecol Evol 23:327–337. https://doi.org/10.1016/j.tree.2008.02.008

Allendorf FW (2013) Conservation and the genetics of populations, 2nd ed. John Wiley & Sons, Hoboken

Araki H, Berejikian BA, Ford MJ, Blouin MS (2008) Fitness of hatchery-reared salmonids in the wild: Fitness of hatchery fish. Evol Appl 1:342–355. https://doi.org/10.1111/j.1752-4571.2008.00026.x

Araki H, Cooper B, Blouin MS (2007a) Genetic effects of captive breeding cause a rapid, cumulative fitness decline in the wild. Science 318:100–103. https://doi.org/10.1126/science.1145621

Araki H, Schmid C (2010) Is hatchery stocking a help or harm? Aquaculture 308:S2–S11. https://doi.org/10.1016/j.aquaculture.2010.05.036

Araki H, Waples RS, Blouin MS (2007b) A potential bias in the temporal method for estimating Ne in admixed populations under natural selection. Mol Ecol 16:2261–2271. https://doi.org/10.1111/j.1365-294X.2007.03307.x

Archer FI, Adams PE, Schneiders BB (2017) STRATAGL: An R package for manipulating, summarizing and analysing population genetic data. Mol Ecol Resour 17:5–11. https://doi.org/10.1111/1755-0998.12559

Balian EV, Segers H, Lévèque C, Martens K (2008) The freshwater animal diversity assessment: an overview of the results. Hydrobiologia 595:627–637. https://doi.org/10.1007/s10750-007-9246-3

BBC (2010) “Perfect storm” caused algae on Llyn Padarn lake. http://news.bbc.co.uk/1/hi/wales/north_west/8616288.stm. Accessed 16 June 2020

Butterworth AJ (1980) The biology of the Arctic char, Salvelinus alpinus L., of Llynnau Peris and Padarn: with special reference to the Dinorwic Reservoir Scheme. Dissertation, University of Liverpool

Caroffino DC, Miller LM, Kapuscinski AR, Ostazeski JJ (2008) Stocking success of local-origin fry and impact of hatchery ancestry: monitoring a new steelhead (Oncorhynchus mykiss) stocking program in a Minnesota tributary to Lake Superior. Can J Fish Aquat Sci 65:309–318. https://doi.org/10.1139/f07-167

Ceballos G, Ehrlich PR, Dirzo R (2017) Biological annihilation via the ongoing sixth mass extinction signaled by vertebrate population losses and declines. PNAS 114:E6089–E6096. https://doi.org/10.1073/pnas.1704949114

Chilcote MW, Goodson KW, Falcy MR (2011) Reduced recruitment performance in natural populations of anadromous salmonids associated with hatchery-reared fish. Can J Fish Aquat Sci 68:511–522. https://doi.org/10.1139/F10-168

Child A (1977) Biochemical polymorphism in char (Salvelinus alpinus L.) from Llynnau Peris, Padarn, Cwellyn and Bodlyn. Heredity 38:359–365

Christie MR, Marine ML, French RA, et al (2012a) Effective size of a wild salmonid population is greatly reduced by hatchery supplementation. Heredity 109:254–260. https://doi.org/10.1038/hdy.2012.39

Christie MR, Marine ML, French RA, Blouin MS (2012b) Genetic adaptation to captivity can occur in a single generation. PNAS 109:238–242. https://doi.org/10.1073/pnas.1111073109

Clabburn P, Davies R, Griffiths J (2014) Summary of the results of hydroacoustic surveys of Llyn Padarn and Llyn Cwellyn, 2013. Natural Resources Wales, Cardiff

DCWW (2015) Drought plan 2015. Dŵr Cymru Welsh Water

Do C, Waples RS, Peel D, et al (2014) NeEstimator v2: re-implementation of software for the estimation of contemporary effective population size (Ne) from genetic data. Mol Ecol Resour 14:209–214. https://doi.org/10.1111/1755-0998.12157

Dray S, Dufour A-B (2007) The ade4 package: implementing the duality diagram for ecologists. J Stat Softw 22:1–20

Elner JK, Happey-Wood CM, Wood DGE (1980) The history of two linked but contrasting lakes in North Wales from a study of pollen, diatoms and chemistry in sediment cores. J Ecolo 68:95–121. https://doi.org/10.2307/2259246

Evanno G, Regnaut S, Goudet J (2005) Detecting the number of clusters of individuals using the software structure: a simulation study. Mol Ecol 14:2611–2620. https://doi.org/10.1111/j.1365-294X.2005.02553.x

Excoffier L, Smouse PE, Quattro JM (1992) Analysis of molecular variance inferred from metric distances among DNA haplotypes: application to human mitochondrial DNA restriction data. Genetics 131:479–491

Ferguson A, Adams C, Jóhannsson M, et al (2019) Trout and char of the North Atlantic Isles. In: Kershner JL, Williams JE, Gresswell RE, Lobón-Cerviá J (eds) Trout and char of the world. American Fisheries Society, Bethesda, Maryland, pp 313–350

Figge F (2004) Bio-folio: Applying portfolio theory to biodiversity. Biodiver Conserv 13:827–849. https://doi.org/10.1023/B:BIOC.0000011729.93889.34

Franklin IR (1980) Evolutionary change in small populations. In: Conservation Biology -An evolutionary-ecological perspective. Sinauer Associates, U.S.A, Sunderland, Massachusetts, pp 135–149

Fraser DJ (2008) How well can captive breeding programs conserve biodiversity? A review of salmonids. Evol Appl 0:080602014503553-??? https://doi.org/10.1111/j.1752-4571.2008.00036.x

Garza JC, Williamson EG (2001) Detection of reduction in population size using data from microsatellite loci. Mol Ecol 10:305–318. https://doi.org/10.1046/j.1365-294X.2001.01190.x

Gomaa NH, Montesinos-Navarro A, Alonso-Blanco C, Picó FX (2011) Temporal variation in genetic diversity and effective population size of Mediterranean and subalpine Arabidopsis thaliana populations. Mol Ecol. https://doi.org/10.1111/j.1365-294X.2011.05193.x

Goudet J (2005) hierfstat, a package for r to compute and test hierarchical F-statistics. Mol Ecol Notes 5:184–186. https://doi.org/10.1111/j.1471-8286.2004.00828.x

Günther A (1862) Contribution to the Knowledge of the British Charrs. In: Proceedings of the Zoological Society of London. pp 37–54

Hatton-Ellis TW (2016) Evidence review of lake nitrate vulnerable zones in Wales. Natural Resources Wales, Bangor

Hauser L, Adcock GJ, Smith PJ, et al (2002) Loss of microsatellite diversity and low effective population size in an overexploited population of New Zealand snapper (Pagrus auratus). PNAS 99:11742–11747

ICES (2019) Working group on North Atlantic salmon. https://doi.org/10.17895/ICES.PUB.4978

IUCN (2019) Global freshwater fish assessment. https://www.iucn.org/theme/species/our-work/freshwater-biodiversity/our-projects/global-freshwater-fish-assessment. Accessed 27 Jun 2020

Jombart T (2008) adegenet: a R package for the multivariate analysis of genetic markers. Bioinformatics 24:1403–1405. https://doi.org/10.1093/bioinformatics/btn129

Keenan K, McGinnity P, Cross TF, et al (2013) diveRsityL: An R package for the estimation and exploration of population genetics parameters and their associated errors. Methods Ecol Evol 4:782–788. https://doi.org/10.1111/2041-210X.12067

Klemetsen A (2010) The charr problem revisited: Exceptional phenotypic plasticity promotes ecological speciation in postglacial lakes. Freshw Rev 3:49–74. https://doi.org/10.4290/FRJ-3.1.3

Kovach R, Jonsson B, Jonsson N, et al (2019) Climate Change and the Future of Trout and Char. In: Kershner JL, Williams JE, Gresswell RE, Lobón-Cerviá J (eds) Trout and char of the world. American Fisheries Society, Bethesda, Maryland, pp 685–716

Lackey RT, Lach D, Duncan S (2006) Salmon 2100: The future of wild Pacific salmon. American Fisheries Society, Bethesda, Md.

Laikre L, Schwartz MK, Waples RS, Ryman N (2010) Compromising genetic diversity in the wild: unmonitored large-scale release of plants and animals. Trends Ecol Evol 25:520–529. https://doi.org/10.1016/j.tree.2010.06.013

Maitland PS, Winfield IJ, McCarthy ID, Igoe F (2007) The status of Arctic charr Salvelinus alpinus in Britain and Ireland. Ecology Freshw Fish 16:6–19. https://doi.org/10.1111/j.1600-0633.2006.00167.x

Mathers RG, De Carlos M, Crowley K, Ó Teangana D (2002) A review of the potential effect of Irish hydroelectric installations on Atlantic salmon (Salmo salar L.) populations, with particular reference to the River Erne. Biol Environ 102:69–79. https://doi.org/10.3318/BIOE.2002.102.2.69

McCarthy ID (2007) The Welsh Torgoch (Salvelinus alpinus): A short review of its distribution and ecology. Ecol Freshw 16:34–40. https://doi.org/10.1111/j.1600-0633.2006.00166.x

Muhlfeld C, Dauwalter D, D’angelo V, et al (2019) Global status of trout and char: Conservation challenges in the twenty-first century. In: Kershner JL, Williams JE, Lobón-Cerviá J (eds) Trout and char of the world. American Fisheries Society, pp 717–760

Nei M (1973) Analysis of gene diversity in subdivided populations. PNAS 70:3321–3323. https://doi.org/10.1073/pnas.70.12.3321

Neville H, Dunham J, Rosenberger A, et al (2009) Influences of wildfire, habitat size, and connectivity on trout in headwater streams revealed by patterns of genetic diversity. Trans Am Fish Soc 138:1314–1327. https://doi.org/10.1577/T08-162.1

Nowak C, Vogt C, Pfenninger M, et al (2009) Rapid genetic erosion in pollutant-exposed experimental chironomid populations. Environ Pollut 157:881–886. https://doi.org/10.1016/j.envpol.2008.11.005

Paradis E (2010) pegas: an R package for population genetics with an integrated-modular approach. Bioinformatics 26:419–420. https://doi.org/10.1093/bioinformatics/btp696

Pauls SU, Nowak C, Bálint M, Pfenninger M (2013) The impact of global climate change on genetic diversity within populations and species. Mol Ecol 22:925–946. https://doi.org/10.1111/mec.12152

Perrier C, Guyomard R, Bagliniere J-L, et al (2013) Changes in the genetic structure of Atlantic salmon populations over four decades reveal substantial impacts of stocking and potential resiliency. Ecol Evol 3:2334–2349. https://doi.org/10.1002/ece3.629

Pritchard JK, Stephens M, Donnelly P (2000) Inference of population structure using multilocus genotype data. Genetics 155:945–959

Quiñones RM, Johnson ML, Moyle PB (2014) Hatchery practices may result in replacement of wild salmonids: adult trends in the Klamath basin, California. Environ Biol Fishes 97:233–246. https://doi.org/10.1007/s10641-013-0146-2

R Core Team (2016) R: A language and environment for statistical computing. R Foundation for Statistical Computing, Vienna, Austria

Reed DH, Frankham R (2003) Correlation between fitness and genetic diversity. Conserv Biol 17:230–237. https://doi.org/10.1046/j.1523-1739.2003.01236.x

Roberts JJ, Fausch KD, Peterson DP, Hooten MB (2013) Fragmentation and thermal risks from climate change interact to affect persistence of native trout in the Colorado River basin. Glob Change Biol 19:1383–1398. https://doi.org/10.1111/gcb.12136

Sgrò CM, Lowe AJ, Hoffmann AA (2011) Building evolutionary resilience for conserving biodiversity under climate change: Conserving biodiversity under climate change. Evol Appl 4:326–337. https://doi.org/10.1111/j.1752-4571.2010.00157.x

Skúlason S, Snorrason SS, Jonsson B (1999) Sympatric morphs, populations and speciation in freshwater fish with emphasis on Arctic charr. In: Magurran AE, May RM (eds) Evolution of biological diversity. Oxford, pp 70–92

Skúlason S, Snorrason SS, Noakes DL, Ferguson MM (1996) Genetic basis of life history variations among sympatric morphs of Arctic char Salvelinus alpinus. Can J Fish Aquat Sci 53:1807–1813

Ursenbacher S, Monney J-C, Fumagalli L (2009) Limited genetic diversity and high differentiation among the remnant adder (Vipera berus) populations in the Swiss and French Jura Mountains. Conserv Genet 10:303–315. https://doi.org/10.1007/s10592-008-9580-7

Valiquette E, Perrier C, Thibault I, Bernatchez L (2014) Loss of genetic integrity in wild lake trout populations following stocking: insights from an exhaustive study of 72 lakes from Québec, Canada. Evol Appl 7:625–644. https://doi.org/10.1111/eva.12160

Waples RS, Do C (2010) Linkage disequilibrium estimates of contemporary N_e_ using highly variable genetic markers: a largely untapped resource for applied conservation and evolution. Evol Appl 3:244–262. https://doi.org/10.1111/j.1752-4571.2009.00104.x

Weeks AR, Sgro CM, Young AG, et al (2011) Assessing the benefits and risks of translocations in changing environments: a genetic perspective: Translocations in changing environments. Evol Appl 4:709–725. https://doi.org/10.1111/j.1752-4571.2011.00192.x

White T (2012) Reasons responsible for the decrease in population size of Llyn Padarn charr (Salvelinus alpinus perisii). Masters thesis

Whiteley AR, Hastings K, Wenburg JK, et al (2010) Genetic variation and effective population size in isolated populations of coastal cutthroat trout. Conserv Genet 11:1929–1943. https://doi.org/10.1007/s10592-010-0083-y

Wickham H (2016) ggplot2: elegant graphics for data analysis, Second edition. Springer, Cham

Wilson AJ, Gíslason D, Skúlason S, et al (2004) Population genetic structure of Arctic charr, Salvelinus alpinus from northwest Europe on large and small spatial scales. Mol Ecol 13:1129–1142. https://doi.org/10.1111/j.1365-294X.2004.02149.x

Winfield IJ, Hateley J, Fletcher JM, et al (2010) Population trends of Arctic charr (Salvelinus alpinus) in the UK: assessing the evidence for a widespread decline in response to climate change. Hydrobiologia 650:55–65. https://doi.org/10.1007/s10750-009-0078-1

