## Appendix for "Genetic insights into the management and conservation of Arctic charr in North Wales"

**Table S1** Multiplex design for Arctic charr in North Wales using 7 microsatellite loci.

| **Locus** | **Primer Sequence (5’-3’)** | **Reference** | **Allele size (bp)** | **Repeat motif** | **Dye** | **Multiplex** | **Primer concentration (uM)** |
| --- | --- | --- | --- | --- | --- | --- | --- |
| Ssa406 | F: AGGTGGGTCCTCCAAGCTAC | O’Reilly et al. (1996) | 404-466 | (GT)14 | NED | 1 | 0.22 |
|  | R: ACCCGCTCCTCACTTAATC |  |  |  |  |  |  |
| Ssa85 | F: AGGTGGGTCCTCCAAGCTAC | O’Reilly et al. (1996) | 147-200 | (GT)14 | NED | 1 | 0.1 |
|  | R: ACCCGCTCCTCACTTAATC |  |  |  |  |  |  |
| SsaD48 | F: GAGCCTGTTCAGAGAAATGAG | King et al. (2005) | 139-300 | TAGA | FAM | 1 | 0.18 |
|  | R: CAGAGGTGTTGAGTCAGAGAAG |  |  |  |  |  |  |
| SalF56 | F: TGCAGTTCCACAATATATCCC | McGowan et al. (2004) | 180-220 | (TG)25 | VIC | 1 | 0.1 |
|  | R: AAGGGCACACTCAGATTTTG |  |  |  |  |  |  |
| SalJ81 | F: CAGCATAATCACTCCCGC | McGowan et al. (2004) | 105-179 | (GT)33 | PET | 1 | 0.1 |
|  | R: GAAAGCTACCTTGCGTGC |  |  |  |  |  |  |
| SalD39 | F: GGGGAGTCTGTGTTAAGTTGG | McGowan et al. (2004) | 194-320 | (GT)3(AC)23 | FAM | 2 | 0.1 |
|  | R: TGAATGGACGTTCCTCTGAC |  |  |  |  |  |  |
| SalE38 | F: CGCCTTGTCATACATTACACC | McGowan et al. (2004) | 95-240 | (AC)44 | PET | 2 | 0.1 |
|  | R: ACGCTACAGAAACAGGAGAAAG |  |  |  |  |  |  |
| SalP61 | F: CACTTATTAACGCCCACTCCC | McGowan et al. (2004) | 100-140 | (CA)18 | VIC | 2 | 0.07 |
|  | R: TTCACAACCACAGGAAAGAACTC |  |  |  |  |  |  |

**References**

King, T. L., Eackles, M. S., & Letcher, B. H. (2005). Microsatellite DNA markers for the study of Atlantic salmon (Salmo salar) kinship, population structure, and mixed-fishery analyses. *Molecular Ecology Notes*, *5*(1), 130–132. https://doi.org/10.1111/j.1471-8286.2005.00860.x

McGowan, C. R., Davidson, E. A., Woram, R. A., Danzmann, R. G., Ferguson, M. M., & Davidson, W. S. (2004). Ten polymorphic microsatellite markers from Arctic charr (*Salvelinus alpinus*): Linkage analysis and amplification in other salmonids. *Animal Genetics*, *35*(6), 479–481. https://doi.org/10.1111/j.1365-2052.2004.01203.x

O’Reilly, P. T., Hamilton, L. C., McConnell, S. K., & Wright, J. M. (1996). Rapid analysis of genetic variation in Atlantic salmon (Salmo salar) by PCR multiplexing of dinucleotide and tetranucleotide microsatellites. *Canadian Journal of Fisheries and Aquatic Sciences*, *53*(10), 2292–2298.

**Table S2** Per locus estimates of several genetic diversity indices in nine populations of Arctic charr in North Wales. Numbers in bold indicate statistically significant values.

| **Locus** | **Population** | **A_R_** | **H_O_** | **H_E_** | **HWE_hom** | **HWE_het** | **Fis** | **HWE (X^2^)** | **HWE (bonf adj.)** |
| --- | --- | --- | --- | --- | --- | --- | --- | --- | --- |
| SalD39 | Bodlyn | 3.89 | 0.21 | 0.57 | NA | **0.000** | **0.65** | 41.65 | **0.000** |
|  | Cowlyd | 10.11 | 0.63 | 0.9 | **0.000** | NA | **0.31** | 145.33 | **0.000** |
|  | Crafnant | 9.94 | 0.67 | 0.88 | NA | **0.000** | **0.26** | 157.92 | **0.000** |
|  | Cwellyn | 8.81 | 0.75 | 0.85 | NA | **0.017** | **0.13** | 111.34 | 0.920 |
|  | Diwaunedd | 7.16 | 0.91 | 0.82 | 0.230 | NA | -0.06 | 29.04 | 1.000 |
|  | Dulyn | 10.31 | 0.74 | 0.86 | NA | **0.001** | 0.15 | 161.49 | 1.000 |
|  | Fynnon Llygwy | 9.18 | 0.57 | 0.88 | NA | **0.000** | **0.37** | 146.82 | **0.000** |
|  | Padarn08 | 9.97 | 0.85 | 0.89 | NA | **0.045** | 0.07 | 83.43 | 0.288 |
|  | Padarn13 | 9.37 | 0.73 | 0.87 | NA | **0.002** | 0.18 | 133.98 | **0.000** |
| SalF56 | Bodlyn | 1.00 | 0 | 0 | NA | NA | NaN | 0.00 | 1.000 |
|  | Cowlyd | 3.43 | 0.67 | 0.58 | **0.002** | NA | **-0.14** | 35.55 | **0.008** |
|  | Crafnant | 4.03 | 0.65 | 0.66 | NA | 0.140 | 0.03 | 15.47 | 0.224 |
|  | Cwellyn | 3.02 | 0.24 | 0.27 | NA | 0.097 | **0.10** | 8.83 | 1.000 |
|  | Diwaunedd | 2.95 | 0.73 | 0.58 | 0.290 | NA | **-0.21** | 3.52 | 1.000 |
|  | Dulyn | 3.16 | 0.31 | 0.55 | NA | **0.000** | **0.44** | 39.84 | **0.008** |
|  | Fynnon Llygwy | 2.60 | 0.65 | 0.51 | **0.033** | NA | **-0.25** | 5.53 | 0.784 |
|  | Padarn08 | 3.38 | 0.62 | 0.55 | NA | **0.041** | -0.09 | 26.68 | **0.016** |
|  | Padarn13 | 3.52 | 0.7 | 0.61 | NA | 0.104 | **-0.14** | 32.27 | **0.000** |
| SalJ81 | Bodlyn | 1.00 | 0 | 0 | NA | NA | NaN | 0.00 | 1.000 |
|  | Cowlyd | 4.54 | 0.62 | 0.67 | 0.545 | 0.556 | 0.08 | 7.52 | 1.000 |
|  | Crafnant | 5.16 | 0.73 | 0.7 | 0.515 | NA | -0.03 | 15.60 | 1.000 |
|  | Cwellyn | 3.53 | 0.53 | 0.51 | 0.607 | NA | **-0.02** | 1.49 | 1.000 |
|  | Diwaunedd | 3.88 | 0.64 | 0.5 | 0.248 | NA | **-0.23** | 2.40 | 1.000 |
|  | Dulyn | 4.76 | 0.6 | 0.73 | NA | **0.008** | 0.20 | 13.19 | 1.000 |
|  | Fynnon Llygwy | 4.99 | 0.52 | 0.61 | NA | **0.002** | 0.15 | 35.54 | 0.088 |
|  | Padarn08 | 3.92 | 1 | 0.68 | **0.001** | NA | **-0.44** | 24.29 | **0.000** |
|  | Padarn13 | 5.19 | 0.55 | 0.67 | NA | **0.009** | **0.20** | 29.89 | 0.536 |
| SalP61 | Bodlyn | 3.61 | 0.59 | 0.59 | NA | **0.014** | 0.02 | 16.93 | 0.632 |
|  | Cowlyd | 4.70 | 0.75 | 0.77 | 0.485 | 0.571 | 0.03 | 9.06 | 1.000 |
|  | Crafnant | 3.19 | 0.67 | 0.62 | 0.312 | NA | **-0.06** | 3.26 | 1.000 |
|  | Cwellyn | 7.77 | 0.73 | 0.84 | NA | **0.014** | **0.15** | 78.73 | 0.160 |
|  | Diwaunedd | 5.50 | 0.82 | 0.77 | 0.719 | NA | -0.02 | 13.95 | 1.000 |
|  | Dulyn | 4.64 | 0.55 | 0.73 | NA | **0.004** | 0.26 | 25.81 | 0.104 |
|  | Fynnon Llygwy | 4.52 | 0.77 | 0.76 | 0.458 | NA | **0.00** | 3.53 | 1.000 |
|  | Padarn08 | 5.48 | 0.76 | 0.73 | NA | 0.117 | -0.02 | 46.61 | **0.000** |
|  | Padarn13 | 4.44 | 0.73 | 0.69 | 0.323 | NA | **-0.04** | 7.01 | 1.000 |
| Ssa406 | Bodlyn | 2.53 | 0.24 | 0.27 | NA | 0.347 | 0.13 | 1.27 | 1.000 |
|  | Cowlyd | 6.78 | 0.87 | 0.82 | 0.849 | 0.515 | **-0.05** | 22.00 | 1.000 |
|  | Crafnant | 4.07 | 0.82 | 0.69 | 0.053 | NA | **-0.17** | 11.86 | 1.000 |
|  | Cwellyn | 5.74 | 0.64 | 0.74 | NA | **0.005** | **0.15** | 58.76 | 0.568 |
|  | Diwaunedd | 4.87 | 0.64 | 0.65 | NA | 0.117 | **0.07** | 13.75 | 1.000 |
|  | Dulyn | 5.79 | 0.79 | 0.82 | NA | 0.292 | **0.05** | 10.38 | 1.000 |
|  | Fynnon Llygwy | 6.92 | 0.76 | 0.81 | NA | **0.005** | 0.08 | 76.09 | 1.000 |
|  | Padarn08 | 6.76 | 1 | 0.83 | **0.023** | NA | **-0.19** | 71.17 | **0.032** |
|  | Padarn13 | 6.29 | 0.7 | 0.78 | NA | 0.088 | 0.12 | 22.26 | 1.000 |
| Ssa85 | Bodlyn | 3.45 | 0.1 | 0.44 | NA | **0.000** | **0.77** | 42.83 | **0.000** |
|  | Cowlyd | 4.45 | 0.7 | 0.67 | **0.030** | 0.661 | -0.02 | 15.67 | 0.216 |
|  | Crafnant | 5.08 | 0.65 | 0.76 | NA | **0.016** | 0.16 | 34.64 | **0.000** |
|  | Cwellyn | 3.71 | 0.49 | 0.56 | NA | **0.005** | 0.15 | 39.82 | 0.808 |
|  | Diwaunedd | 2.00 | 0.55 | 0.5 | 0.671 | NA | **-0.05** | 0.11 | 1.000 |
|  | Dulyn | 4.66 | 0.58 | 0.66 | NA | 0.085 | 0.14 | 25.29 | 0.592 |
|  | Fynnon Llygwy | 4.22 | 0.5 | 0.54 | NA | 0.180 | 0.08 | 17.04 | 1.000 |
|  | Padarn08 | 6.36 | 0.43 | 0.75 | NA | **0.001** | **0.45** | 60.52 | **0.000** |
|  | Padarn13 | 4.32 | 0.55 | 0.63 | NA | 0.121 | 0.14 | 26.63 | 0.160 |
| SsaD48 | Bodlyn | 7.64 | 0.93 | 0.82 | 0.084 | NA | **-0.12** | 92.09 | 1.000 |
|  | Cowlyd | 7.81 | 0.77 | 0.79 | 0.894 | NA | 0.04 | 45.14 | 1.000 |
|  | Crafnant | 9.37 | 0.54 | 0.82 | NA | **0.000** | **0.35** | 331.96 | **0.000** |
|  | Cwellyn | 8.93 | 0.92 | 0.87 | 0.276 | NA | **-0.04** | 130.25 | 1.000 |
|  | Diwaunedd | 6.39 | 0.82 | 0.78 | 0.461 | NA | **-0.01** | 35.38 | 1.000 |
|  | Dulyn | 7.62 | 0.77 | 0.75 | 0.279 | NA | **-0.01** | 71.74 | 1.000 |
|  | Fynnon Llygwy | 7.91 | 0.74 | 0.8 | NA | **0.050** | 0.08 | 82.47 | 0.680 |
|  | Padarn08 | 8.82 | 0.95 | 0.85 | 0.283 | NA | -0.09 | 149.52 | **0.000** |
|  | Padarn13 | 8.52 | 0.82 | 0.76 | 0.224 | NA | **-0.06** | 69.19 | 1.000 |

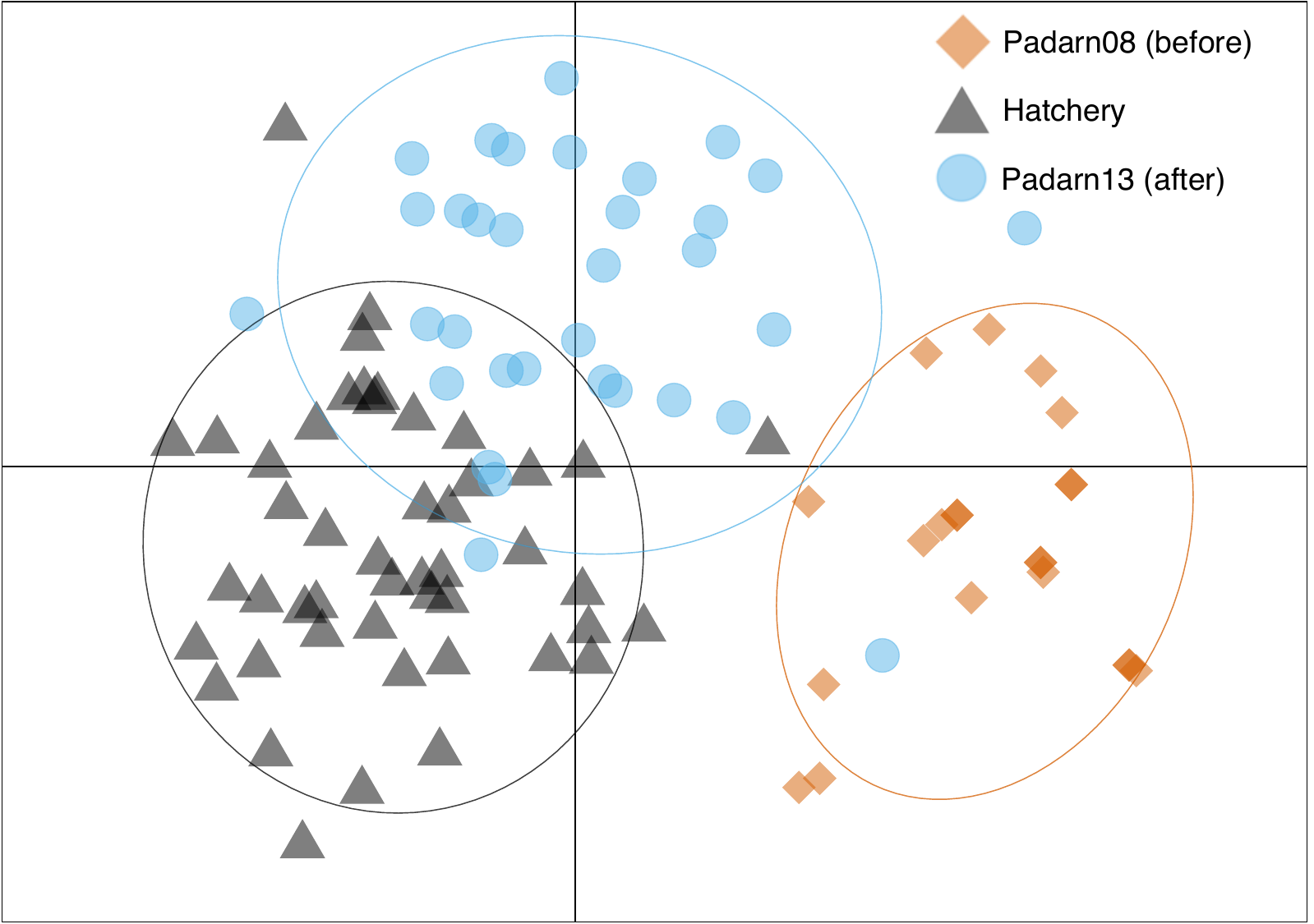

**Figure S1** Discriminate Analysis of Principal Components (DACP) for Llyn Padarn before (Padarn08) and after (Padarn13) hatchery supplementation from Crafnant.
